# Altered MicroRNA Signatures in Circulating Extracellular Vesicles Reflect Aortic Dilation in Takayasu Arteritis

**DOI:** 10.1101/2025.09.21.677656

**Authors:** Kohei Kawajiri, Shun Nakagama, Tetsuo Sasano, Yasuhiro Maejima

## Abstract

**Background:** Takayasu arteritis (TAK) is a large-vessel vasculitis characterized by progressive inflammation that can lead to stenosis or aneurysmal dilation. Reliable circulating biomarkers for vascular remodeling are lacking, and extracellular vesicles (EVs), which transport molecular cargo that reflects their cellular origin, have emerged as potential biomarkers and modulators of vascular inflammation. However, their role in TAK remains unclear. This study aimed to investigate the relationship between EV-derived microRNA (miRNA) signatures and clinically relevant vascular phenotypes, with a primary focus on aortic dilation.

**Methods:** Circulating small EVs (sEVs) were isolated from the serum of patients with TAK and healthy controls, characterized by high-sensitivity flow cytometry, were profiled for miRNA content, and analyzed using bioinformatic pathway enrichment. In vitro assays were employed in assessing their impact on endothelial activation, while the diagnostic performance of miRNAs was compared to conventional biomarkers and clinical parameters.

**Results:** sEVs from patients with TAK upregulated the endothelial expression of intercellular adhesion molecule-1 and vascular cell adhesion molecule-1, thereby enhancing monocyte adhesion in vitro, consistent with a pro-inflammatory phenotype. Differentially expressed miRNAs were enriched in PI3K–Akt signaling, with phosphatase and tensin homolog identified as a key predicted target. Among patients, miR-223-3p was selectively downregulated in those with aortic dilation, regardless of disease activity or treatment status. Receiver operating characteristic analysis showed that miR-223-3p identified aortic dilation with an area under the curve of 0.748, outperforming C-reactive protein and erythrocyte sedimentation rate. Flow cytometric analysis suggested that platelet-derived EVs represent a predominant source of circulating miR-223-3p, and their proportion tended to be reduced in patients with aortic dilation.

**Conclusions:** Reduced platelet-derived EV–associated miR-223-3ps are associated with aortic dilation in TAK and demonstrate superior diagnostic accuracy over conventional inflammatory markers. EV miRNA profiling may provide a novel approach for evaluating vascular remodeling in TAK.

## Introduction

Takayasu arteritis (TAK) is an idiopathic large-vessel vasculitis that primarily involves the aorta and its major branches, resulting in progressive vascular stenosis, occlusion, or aneurysmal dilation of the aorta^1^, and it is most prevalent among young and middle-aged women of Asian descent^2–4^. Despite advances in imaging and the development of immunosuppressive therapies, early diagnosis of TAK remains challenging, and the mechanisms driving vascular remodeling and disease progression remain partially understood. Reliable circulating biomarkers that reflect disease activity, vascular injury, or progression toward aortic dilatation are urgently needed to improve clinical management. Several serum biomarkers have been proposed in TAK, including pentraxin-3 (PTX3)^5^, apolipoprotein C-2^6^, serum amyloid A^7^, interleukin (IL)-6^8^, IL-12^9^, IL-18^10^, matrix metalloproteinases^11^, and soluble intercellular adhesion molecule-1 (ICAM-1)^12^. However, in clinical practice, none have demonstrated superiority over conventional biomarkers, such as C-reactive protein (CRP) or erythrocyte sedimentation rate (ESR), for monitoring disease activity^13–15^.

Extracellular vesicles (EVs) are nanosized, phospholipid bilayer–enclosed particles that transport bioactive molecules such as proteins, lipids, and nucleic acids, thereby mediating intercellular communication^16, 17^. Increasing evidence suggests that EVs are increasingly recognized as critical modulators of vascular inflammation, endothelial dysfunction^18^, and immune cell activation^19^. Notably, the molecular cargo of EVs—particularly microRNAs (miRNAs)—reflects the activation state and cellular origin of their parent cells, highlighting their potential as minimally invasive biomarkers^17^. Nonetheless, in TAK, the pathogenic role and biomarker utility of EV-associated miRNAs remain unclear.

This study aimed to investigate the relationship between EV-derived miRNA signatures and clinically relevant vascular phenotypes, with a primary focus on aortic dilation.

## Methods

### Patients and Sample Collection

This study was approved by the Ethics Committee on Genetic Research of the School of Medicine, Institute of Science, Tokyo (Approval No. G2000-180). A total of 54 patients diagnosed with TAK, who presented to the Institute of Science Tokyo Hospital and fulfilled the 1990 American College of Rheumatology (ACR) classification criteria for TAK^3^, were enrolled. Serum samples were collected from both patients with TAK and healthy volunteers. Blood samples were obtained from patients with TAK during routine clinical care, and clinical data were retrospectively collected by physicians. Serum samples were stored at –80°C at the Institute of Science Tokyo Bioresource Research Support Center until analysis. Patients were classified according to the angiographic classification proposed at the International TAK Conference in Tokyo in 1994 (Numano’s classification)^20^, which categorizes vascular lesions as follows: Types I (branches of the aortic arch), IIa (ascending aorta, aortic arch, and its branches), IIb (ascending aorta, aortic arch and its branches, and thoracic descending aorta), III (thoracic descending aorta, abdominal aorta, and/or renal arteries), IV (abdominal aorta and/or renal arteries), and V (combined features of Types IIb and IV).

Disease activity was assessed using the criteria proposed by Kerr et al.^21^, with patients meeting at least two of the four criteria being classified as having active disease. For the assessment of arterial dilation, nine aortic regions, including their branches, as defined in the 2022 ACR and European Alliance of Associations for Rheumatology Classification Criteria for Takayasu Arteritis (imaging domain)^22^, were evaluated. Either radiologists or cardiologists determined arterial dilation based on imaging findings from computed tomography (CT) angiography, magnetic resonance imaging (MRI), positron emission tomography (PET)-CT, or vascular ultrasonography.

### Isolation of Serum Small EVs (sEVs) in Humans

Isolation of sEV was based on the recent position statement of the International Society for Extracellular Vesicles (MISEV, 2023)^23^. Venous blood was collected and centrifuged at 1,700 ×g for 10 min at 4°C to separate the serum, which was immediately stored at −80°C until further use. In each experiment, samples were thawed and subjected to sEV isolation using size exclusion chromatography (SEC). sEVs were isolated with the qEVoriginal GEN2 70 nm columns (Izon Science, Oxford, UK) following the manufacturer’s protocol. In summary, serum samples were first centrifuged at 1,500 ×g for 10 min to remove cells and large particles. The supernatant was collected and centrifuged at 10,000 ×g for another 10 min. Subsequently, 500 µL of the resulting supernatant was loaded onto the pre-equilibrated SEC column. Elution was carried out using phosphate-buffered saline (PBS) that had been filtered through a 0.22 μm membrane, and 16 sequential fractions of 400 µL each were collected. Among these, fractions 6–16 were used for the assessment of particle number, size distribution, and purity. Particle size and concentration were measured using interferometric light microscopy (ILM) with the Videodrop system (Myriade, France). Protein concentrations were determined using the Pierce bicinchoninic (BCA) Protein Assay Kit (Thermo Fisher Scientific), and the purity of each fraction was calculated as the ratio of particle number to protein content. For the verification of successful isolation of sEV, immunoblotting was performed for specific sEV markers (CD63 and TSG101), with apoliprotein A-1 (ApoA1) and ATP5A as negative controls (NCs). In addition, the morphology of isolated sEVs was assessed using transmission electron microscopy (TEM) (JEM-1400Flash, JEOL Ltd.), and the isolated sEVs were aliquoted and stored at −80°C, using a single aliquot for each subsequent experiment.

### Cell Lines and Cell Culture

EA.hy926 cells were cultured in Dulbecco’s Modified Eagle’s Medium (DMEM; Nacalai Tesque, Japan) supplemented with 10% fetal bovine serum (FBS) and 1% penicillin-streptomycin (Nacalai Tesque). THP-1 cells were maintained in RPMI-1640 medium (Nacalai Tesque) containing 10% FBS and 1% penicillin-streptomycin. All cells were incubated at 37°C in a humidified atmosphere with 5% carbon dioxide (CO₂). For selected experiments, THP-1 cells were differentiated with phorbol 12-myristate 13-acetate (PMA) (AdipoGen) and subsequently stimulated with lipopolysaccharide (LPS) (Invitrogen). For stimulation experiments, sEVs were added at a final concentration of 2.0×10^9^ particles/mL.

### Adhesion Assay of THP-1 to Endothelial Cells

To assess the adhesion of THP-1 cells to endothelial cells (ECs), THP-1 cells were labeled with the fluorescent dye calcein-AM (FUJIFILM Wako Pure Chemical Corporation, Japan). Cells were incubated in RPMI-1640 medium supplemented with 10% FBS and 10 μM calcein-AM at 37°C for 1 h. Dye loading was terminated by adding cold RPMI-1640 containing 10% FBS, followed by centrifugation. The fluorescence-labeled cells were resuspended in RPMI-1640 supplemented with 10% FBS and used for adhesion assays.

ECs were cultured in 35-mm glass-bottom dishes (Matsunami Glass, Japan) until they attained confluence. Monolayers were stimulated for 24 h with PBS, healthy control (HC)-derived EVs, or Takayasu arteritis-derived EVs (TAK-EVs). Thereafter, 1 × 10^5^ calcein-AM-labeled THP-1 cells were added to each dish and co-cultured for 1 h in the absence of light at 37°C under 5% CO₂ with gentle shaking. Non-adherent THP-1 cells were removed by rinsing the monolayer with PBS. The number of fluorescently labeled monocytes adhering to the EC surface was evaluated using a fluorescence microscope (Keyence, Japan). Five random fields per dish were selected, and the adherent monocytes were manually counted.

### RNA Extraction from Cultured Cells and Quantitative Polymerase Chain Reaction (qPCR)

Total RNA was extracted from cultured cells using the RNeasy Mini Kit (QIAGEN, Hilden, Germany) following the manufacturer’s instructions. Complementary DNA (cDNA) was synthesized from 200 ng of total RNA with the High-Capacity cDNA Reverse Transcription Kit (Thermo Fisher Scientific, Waltham, MA, USA).

qPCR was performed on a StepOnePlus Real-Time PCR System (Thermo Fisher Scientific) using Power SYBR Green Master Mix (Thermo Fisher Scientific) and custom-designed primers. Gene expression was normalized to *Gapdh* (glyceraldehyde-3-phosphate dehydrogenase), and relative expression was calculated using the ΔΔCt method. Primer sequences are listed in Table S1.

### Immunoblotting

Protein extracts from THP-1 cells were prepared using radioimmunoprecipitation assay buffer (Nacalai Tesque). Protein concentrations were determined using the BCA Protein Assay Kit. Equal amounts of protein (10 µg per lane) were separated by sodium dodecyl sulphate-polyacrylamide gel electrophoresis (TGX FastCast, Bio-Rad Laboratories, Hercules, CA, USA) and transferred to polyvinylidene fluoride membranes using the Trans-Blot Turbo transfer system (Bio-Rad). Following blocking with 5% skim milk in tris-buffered saline, the membranes were incubated overnight at 4°C with primary antibodies: mouse anti-CD63 (Santa Cruz, sc-5275; 1:500), mouse anti-TSG101 (Santa Cruz, sc-7964; 1:500), rabbit anti-APOA1 (Abcam ab52945; 1:500), mouse anti-ATP5A (Abcam ab14748; 1:500), rabbit anti-NLR family pyrin domain containing 3 (NLRP3) (Cell signaling, 15101S; 1:1000), and mouse anti-GAPDH (Santa Cruz, sc-32233; 1:10000). This was followed by incubation with horseradish peroxidase conjugated secondary antibodies: Anti-mouse immunoglobulin (IgG), HRP-linked Antibody (Cell Signaling, 7076S; 1:10000), Anti-rabbit IgG, and HRP-linked Antibody (Cell Signaling, 7074S; 1:10000). Chemiluminescent signals were detected using the iBright CL1500 imaging system (Thermo Fisher Scientific). Band intensities were quantified with ImageJ software and normalized to GAPDH.

### miRNA Microarray

Total RNA was extracted from fractions seven and eight of serum-derived sEVs, which were isolated by the aforementioned SEC method, using the 3D-Gene RNA extraction reagent (Toray Industries, Tokyo, Japan). Comprehensive miRNA expression profiling was performed using the 3D-Gene miRNA Labeling Kit and the 3D-Gene Human miRNA Oligo Chip (Toray Industries), designed to detect all miRNA sequences registered in miRBase (http://www.mirbase.org/).

Global normalization was applied by adjusting the median of the total signal intensity for each sample to a value of 25. The presence of each miRNA was defined as a microarray signal exceeding the threshold, defined as the mean plus two standard deviations (SD) of the NC signals. After a miRNA was considered present, the mean of the NC signals (after removal of the top and bottom 5% of values based on intensity) was subtracted from the corresponding miRNA signal. Whenever the background-subtracted signal was negative or undetectable, a value of 1 was assigned on a linear scale. In addition, miRNAs that were expressed below the detection limit (low signal) in more than 50% of samples within either group were excluded from further analysis. All microarray data generated in this study complied with the Minimum Information about Microarray Experiment (MIAME) guidelines^24^.

### miRNA Analysis

Normalized microarray data were used for downstream analyses. Principal component analysis was performed to visualize global patterns of miRNA expression and assess sample clustering. For differential expression analysis, linear models were fitted to each gene using the *lmFit* function from the limma R package (version 3.54.0; https://bioconductor.org/packages/release/bioc/html/limma.html). Empirical Bayes moderation was then applied using the eBayes function to improve the precision of gene-wise variance estimates by borrowing strength across all genes.

For comparisons between HC and patients with TAK, differentially expressed miRNAs were defined as those with a false discovery rate (q-value) < 0.05 and an absolute fold change ≥ 2. For subgroup comparisons within TAK (non-dilated vs. dilated aorta), significance was defined as *p* < 0.05 with a fold change ≥ 2.

To explore the biological relevance of the dysregulated miRNAs, pathway enrichment analysis of their predicted target genes was performed with the DIANA-miRPath v4.0 platform, which allows for the identification of signaling pathways targeted by single or multiple miRNAs^25^. In addition, miRNA–target interactions were further examined with miRTargetLink 2.0^26^, a tool that constructs hierarchical miRNA–gene interaction networks based on experimentally validated and predicted data.

### miRNA Extraction and qPCR Analysis

Total miRNA was extracted from purified sEVs using the miRNeasy Mini Kit (QIAGEN, Hilden, Germany) following the manufacturer’s instructions. cDNA was synthesized from the isolated RNA using the TaqMan™ Advanced miRNA cDNA Synthesis Kit (Thermo Fisher Scientific). qPCR was then performed using the TaqMan™ Fast Advanced Master Mix on a StepOnePlus™ Real-Time PCR System (Thermo Fisher Scientific). Cycle threshold (Ct) values were normalized to the spike-in control ath-miR159a, which was added during the RNA extraction process to correct for technical variation.

### miRNA Mimic Transfection

THP-1 cells were transfected with 50 nM miRNA mimic using Lipofectamine™ RNAiMAX (Invitrogen, Thermo Fisher Scientific) following the manufacturer’s instructions. After 6 h, the medium was replaced with a fresh complete medium, and cells were cultured for 18 h to allow recovery. Subsequently, cells were stimulated with LPS, followed by protein expression analysis.

### Flow Cytometric Analysis for EV Characterization

Serum-derived sEVs were characterized by high-sensitivity flow cytometry using the SA3800 system (Sony, Japan). sEV samples were diluted in 0.22 µm-filtered, calcium- and magnesium-free PBS and incubated with fluorophore-conjugated antibodies for 2 h in the dark. After incubation, samples were washed with PBS (-) using a 300 kDa molecular weight cutoff filter (Vivaspin 500, Sartorius, Germany) and then resuspended in PBS (-) for analysis.

Flow cytometric analysis was performed on the SA3800 system. Controls included unstained and isotype control samples. Gating for particle size was guided by reference beads (Megamix Plus FSC and Megamix Plus SSC; Biocytex, France), and sEVs were analyzed within a size range corresponding to approximately 100–200 nm. The acquired data were analyzed using FlowJo software (version 10; BD Biosciences, USA). Representative gating strategies and antibody staining profiles are presented in Figure S5.

### Statistical Analyses

All statistical analyses were performed using GraphPad Prism version 10 (GraphPad Software, Inc., San Diego, CA) and R version 4.3.1 with appropriate packages. For comparisons between three or more groups, one-way analysis of variance followed by Tukey’s post hoc test was applied whenever the data met assumptions of normality and homogeneity of variance. For comparisons between two groups, unpaired t-tests were used under the same assumptions. Whenever the data were non-normally distributed or the sample size was small (n < 6), and normality could not be reliably assessed, non-parametric tests were used; specifically, the Mann–Whitney U test was applied for two-group comparisons.

Data are presented as mean ± SD, mean ± standard error of the mean (SEM), or median (interquartile range [IQR]: 25th‒75th percentile) as indicated in the figure legends. All experiments were independently repeated at least three times, except for the miRNA transcriptomic analysis. Receiver operating characteristic (ROC) curves were generated, and the area under the curve (AUC) was calculated, with values of 1.0 and 0.5 representing perfect and non-informative discrimination, respectively.

## Results

### Patient Characteristics

A total of 54 patients with TAK, who had been followed at our institution, were included in this study (Table 1). The median age at analysis was 55 years (IQR, 37.5–67.0), and the median age at disease onset was 26 years (IQR, 21–35). According to the Numano classification^20^, nine patients (16.7%) were classified as type I, 21 (38.9%) as type IIa, 12 (22.2%) as type IIb, none as type III, three (5.6%) as type IV, and nine (16.7%) as type V. Surgical vascular interventions had been performed in 11 patients (20.4%). At the time of assessment, seven patients (13.0%) were classified as having active disease based on Kerr’s criteria^21^.

**Table 1.** Clinical characteristics of patients with Takayasu arteritis **Abbreviations**: DMARDs, disease-modifying antirheumatic drugs; Biologics, biologic agents; ESR, erythrocyte sedimentation rate; CRP, C-reactive protein; WBC, white blood cell count.

| Table. Clinical Characteristics of TAK patients |  |
| --- | --- |
| Characteristics | TAK patients (n = 54) |
| Female | 53 (98.1) |
| Age at onset, y | 26 (14) |
| Numano classification |  |
| I | 9 (16.7) |
| II a | 21 (38.9) |
| II b | 12 (22.2) |
| III | 0 |
| IV | 3 (5.6) |
| V | 9 (16.7) |
| Arterial dilation | 12 (22.2) |
| Surgery | 11 (20.4) |
| Active disease | 7 (13.0) |
| Glucocorticoid | 35 (64.8) |
| DMARDs | 3 (5.6) |
| Biologics | 9 (16.7) |
| ESR, mm/h | 13 (22) |
| CRP, mg/dL | 0.08 (0.325) |
| WBC, /μL | 6,700 (3,375) |
| n (%), median (IQR) |  |

### Characterization of Purified Circulating sEVs from Patients with and without TAK

SEC-based isolation of serum from patients with TAK and HC revealed that sEVs were predominantly recovered in fraction 6–16, as confirmed by ILM and BCA assay (Figure 1A). Particle concentrations peaked in fractions seven and eight, whereas protein content increased in later fractions, indicating the presence of non-vesicular protein contamination. Purity, defined as the particle-to-protein ratio, was highest in fractions seven and eight (Figure 1B), which were selected for downstream analyses.

**Figure 1.**
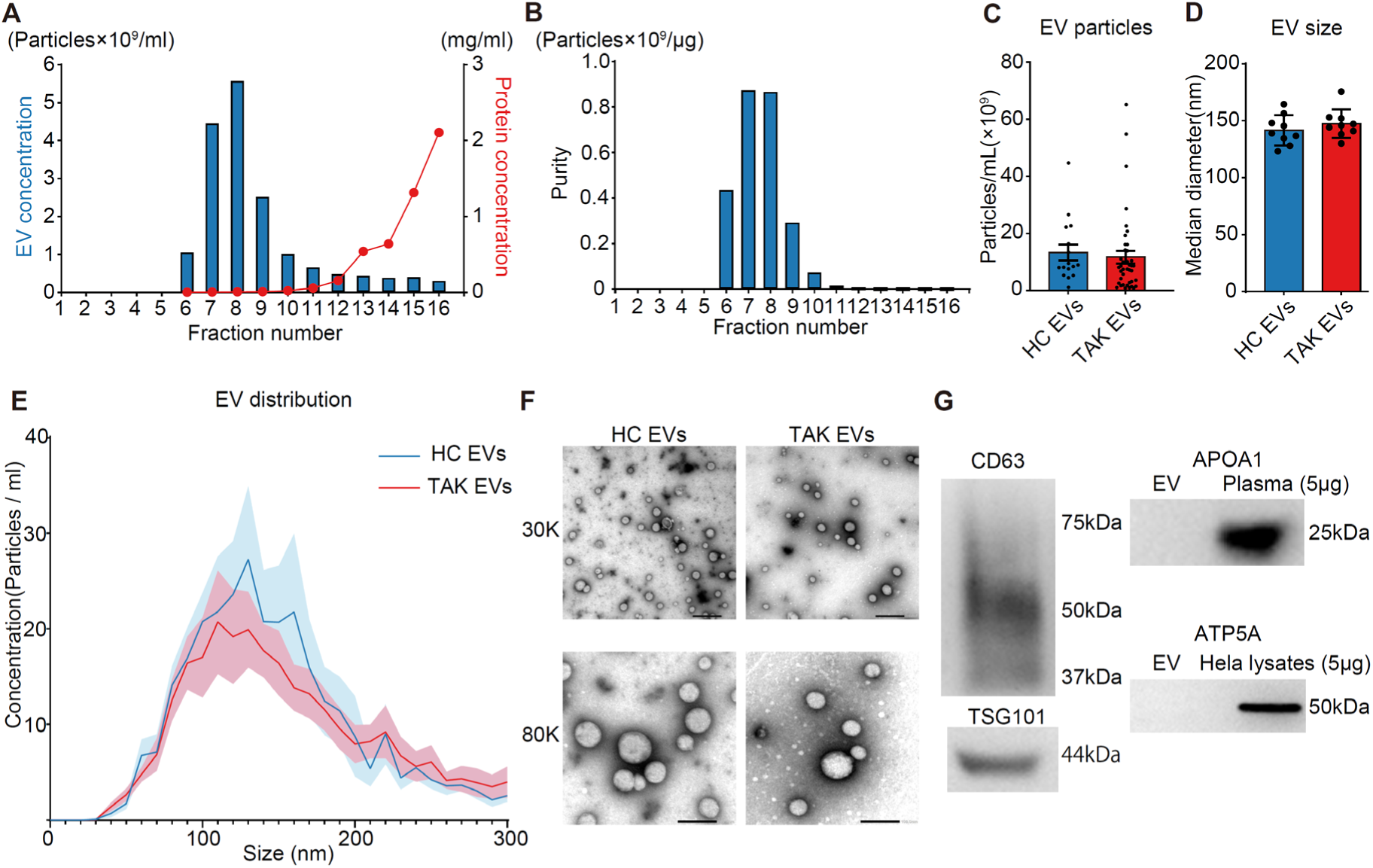
Characterization of circulating sEVs isolated from the serum of patients with TAK and HC (A) Particle concentration and protein content across SEC fractions (fractions 6–16) measured by ILM and BCA assay, respectively. (B) Particle-to-protein ratio for each fraction as an indicator of sEV purity. (C, D) ILM-based measurement of particle concentration (C) and median particle size (D) in sEV preparations from TAK and HC serum. (E) Particle size distribution profiles derived from ILM data. Solid lines represent mean values for TAK (red line, n = 6) and HC (blue line, n = 6); shaded areas indicate SD. (F) Representative TEM images of serum-derived sEVs. The upper image was acquired at 30,000× magnification (scale bar: 200 nm) and the lower image at 80,000× (scale bar: 100 nm). (G) Representative western blot images showing expression of canonical sEV markers (CD63, TSG101) and non-vesicular proteins (ApoA1, ATP5A). Data are presented as mean ± SD or SEM. (C and D) unpaired t-test; *p* < 0.05. **Abbreviations:** TAK, Takayasu arteritis; HC, healthy control; sEV, small extracellular vesicle; SEC, size-exclusion chromatography; ILM, interferometric light microscopy; BCA, bicinchoninic acid; TEM, transmission electron microscopy; TSG101, tumor susceptibility gene 101; ApoA1, apolipoprotein A1; ATP5A, ATP synthase F1 subunit alpha; SD, standard deviation; SEM, standard error of the mean

Comparison of EV concentration and size between TAK and HC groups using ILM revealed no significant differences in either particle concentration or mean size (Figures 1C, D). Size distribution profiles were also similar, with most particles ranging from 50–150 nm in both groups (Figure 1E). TEM confirmed the presence of spherical vesicles within this range (Figure 1F). Western blot analysis validated the enrichment of sEVs, demonstrating robust signals for CD63 and TSG101, established sEV markers, and absence of non-vesicular contaminants such as ApoA1 and ATP5A, thereby supporting the specificity and purity of the isolated sEVs.

Collectively, these findings demonstrate that our SEC-based isolation method reliably yields highly pure, morphologically intact sEVs suitable for downstream analyses.

### Effect of TAK-EVs on Endothelial Activation and Monocyte Adhesion

To explore the cellular targets of sEVs derived from patients with TAK (TAK-EVs), we first examined their effects on macrophages. THP-1 monocytes were differentiated into macrophage-like cells using PMA, followed by stimulation with LPS in the presence of TAK-EVs or HC-EVs. The mRNA levels of key proinflammatory cytokines (tumor necrosis factor-alpha [*TNF-α], IL-6, and IL-1β*) and chemotaxis-related genes (*CCL2, CXCR4, and CXCL1*) were assessed by qPCR (Figure S1). However, no significant differences in gene expression were observed between the TAK and HC groups under these conditions, suggesting that TAK-EVs do not affect LPS-induced activation of macrophages.

To evaluate whether TAK-EVs influence vascular cells, the impact of TAK-EVs on human ECs was assessed. A monocyte adhesion assay using THP-1 cells and ECs revealed that ECs pretreated with TAK-EVs exhibited a significantly increased number of THP-1 adhesion compared to those treated with HC-EVs (Figures 2A, 2B).

**Figure 2.**
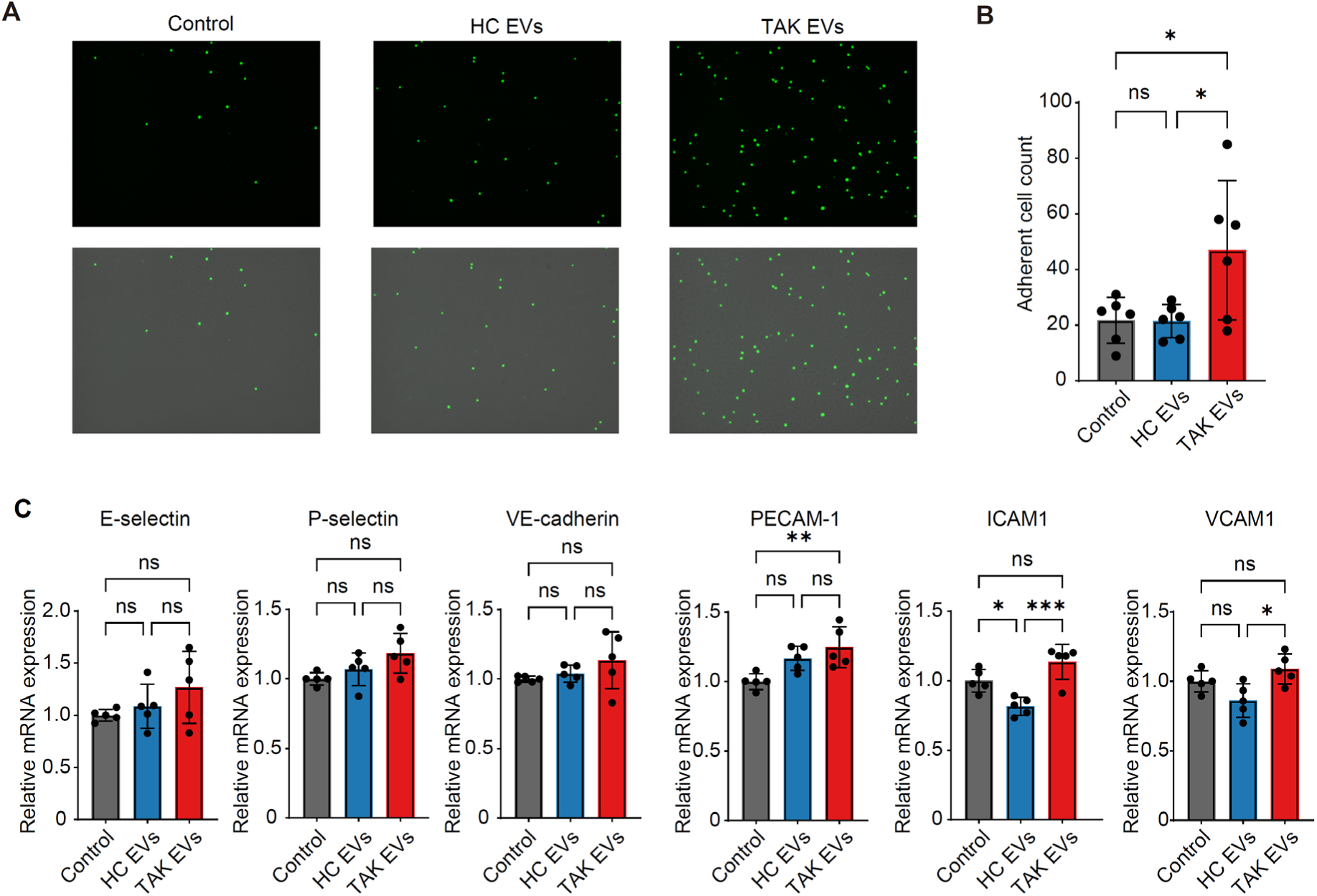
TAK-EVs enhance monocyte adhesion to endothelial cells and upregulate adhesion molecule expression (A) Schematic representation of the monocyte adhesion assay. Human ECs were pretreated with either PBS (left), HC-EVs (middle), or TAK-EVs (right) for 24 h, followed by incubation with calcein-AM-labeled THP-1 monocytes. Representative fluorescence images of adherent THP-1 cells are shown in the upper panel; merged images with underlying ECs are shown in the lower panel. (B) Quantification of adherent THP-1 cells per microscopic field. (C) Relative mRNA expression levels of adhesion-related genes in ECs following treatment with HC-EVs or TAK-EVs, assessed by quantitative PCR. (B and C) Data are presented as mean ± SD, one-way ANOVA followed by Tukey’s post hoc test. (B) **Abbreviations:** EC, endothelial cell; E-selectin, endothelial selectin; P-selectin, platelet selectin; VE-cadherin, vascular endothelial cadherin; PECAM-1, platelet endothelial cell adhesion molecule 1; ICAM-1, intercellular adhesion molecule 1; VCAM-1, vascular cell adhesion molecule 1; SD, standard deviation; SEM, standard error of the mean; PBS, phosphate-buffered saline; ANOVA, analysis of variance; HC-EVs, healthy control–derived extracellular vesicles; TAK-EVs, Takayasu arteritis–derived extracellular vesicles; THP-1, human monocytic cell line; PCR, polymerase chain reaction; mRNA, messenger ribonucleic acid

To further elucidate the underlying mechanisms, the expression of adhesion-related genes in ECs following sEV exposure was analyzed. TAK-EV treatment significantly upregulated the expression of *ICAM-1* and vascular cell adhesion molecule-1 (*VCAM-1*), as determined by qPCR (Figure 2C). These results suggest that TAK-EVs enhance endothelial activation and promote inflammatory cell recruitment, potentially contributing to vascular inflammation and dysfunction in TAK.

### Differential Serum Exosomal miRNA Profiles in Patients with TAK

To investigate the differential expression of miRNAs encapsulated in circulating sEVs, we performed comprehensive miRNA profiling on serum-derived sEVs from patients with TAK and HC. Before profiling, the integrity and enrichment of small RNAs within serum-derived sEVs were confirmed using Bioanalyzer analysis (Figure S2). Principal component analysis clearly separated TAK and HC groups, demonstrating distinct miRNA expression patterns (Figure 3A).

**Figure 3.**
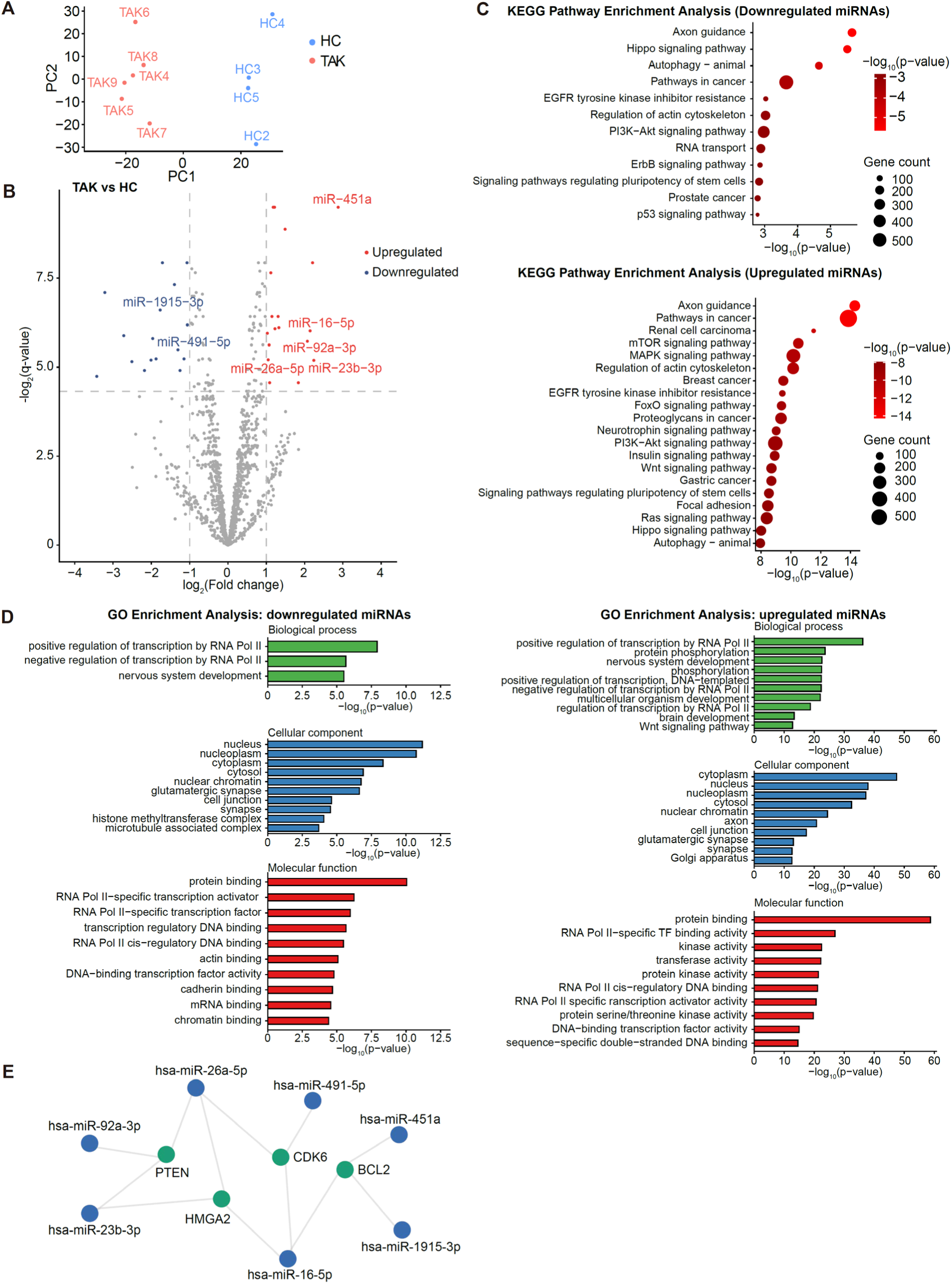
Circulating sEVs from patients with TAK exhibit distinct miRNA profiles and pathway associations (A) PCA of miRNA expression profiles in serum-derived sEVs from patients with TAK and HC. (B) Volcano plot illustrating differential expression of miRNAs between TAK and HC groups. miRNAs with fold change ≥ 2 and q-value < 0.05 were considered significant. (C) Pathway enrichment analysis performed using DIANA-miRPath v4.0 based on the differentially expressed miRNAs. The upper panel represents pathways enriched among downregulated miRNAs, and the lower panel shows pathways enriched among upregulated miRNAs. (D) GO enrichment analysis categorized into three domains: biological processes, cellular components, and molecular functions. GO enriched in downregulated (left) and upregulated (right) miRNAs. (E) miRNA–target gene interaction network constructed using miRTargetLink 2.0 based on both predicted and validated interactions. **Abbreviations:** PCA, principal component analysis; GO, Gene Ontology; sEVs, small extracellular vesicles; TAK, Takayasu arteritis; miRNA, microRNA; HC, healthy control.

Differential expression analysis identified 35 dysregulated miRNAs in TAK-EVs compared to HC-EVs (18 upregulated, 17 downregulated; Fold change ≥ 2, q-value < 0.05), as visualized in the volcano plot (Figure 3B). Pathway enrichment analysis using DIANA-miRPath v4.0^25^ revealed that upregulated miRNAs were predominantly associated with signaling pathways such as mTOR, MAPK, PI3K-Akt, and Wnt signaling, whereas downregulated miRNAs were enriched in Hippo, autophagy, PI3K-Akt, FoxO, and p53 signaling pathways (Figure 3C). These enriched pathways highlight potential contributions of TAK-EV miRNAs to processes including cell proliferation, apoptosis, and autophagy, as well as stress responses. Gene Ontology enrichment analysis showed that upregulated miRNAs were enriched in biological processes such as protein phosphorylation, regulation of transcription by RNA polymerase II, and multicellular organism development (Figure 3D).

Downregulated miRNAs were enriched in processes related to both positive and negative regulation of RNA polymerase II transcription and nervous system development. Enrichment in molecular functions, such as DNA-binding transcription factor activity and protein kinase activity, as well as cellular components like nuclear chromatin and synapses, was also observed (Figure 3D). These findings suggest that TAK-associated miRNAs may modulate transcriptional programs and signaling relevant to immune activation and vascular remodeling.

Finally, miRNA-target interaction network analysis using miRTargetLink 2.0^26^, which prioritizes interactions where individual genes are regulated by multiple miRNAs, identified regulatory hubs in which multiple dysregulated miRNAs converged on common targets (Figure 3E). Notably, *PTEN* (phosphatase and tensin homolog), *HMGA2* (high mobility group AT-hook 2), *CDK6* (cyclin-dependent kinase 6), and *BCL2* (B-cell lymphoma 2) were predicted as central nodes. Among these, *PTEN* is particularly relevant because of its established role in vascular barrier regulation and angiogenesis via PI3K/Akt signaling. These upregulated miRNAs in TAK-EVs, including hsa-miR-23b-3p, hsa-miR-26a-5p, and hsa-miR-92a-3p, were predicted to target PTEN at 3′ untranslated region (3′ UTR) sites, suggesting a cooperative post-transcriptional repression mechanism that could exacerbate vascular inflammation (Figure S3).

In summary, these findings demonstrate that circulating sEVs from patients with TAK harbor a distinct miRNA signature that targets key signaling pathways, with potential consequences for vascular inflammation, endothelial dysfunction, and pathological remodeling.

### Identification of MiRNAs Characteristic of Aortic Dilation in Patients with TAK

Building on the distinct sEV-encapsulated miRNA profiles identified in patients with TAK, the potential association between specific miRNAs and vascular complications, particularly aortic dilation, was assessed. Aortic dilation is a major manifestation of disease progression in advanced TAK^27, 28^; however, its molecular determinants remain poorly understood. To explore potential biomarkers, we compared serum sEV miRNA expression profiles between patients with TAK, with aortic dilation (n = 3), and without dilation (n = 3). Volcano plot analysis identified miR-223-3p as the only significantly differentially expressed miRNA, showing marked downregulation in patients with aortic dilation (Figure 4A).

**Figure 4.**
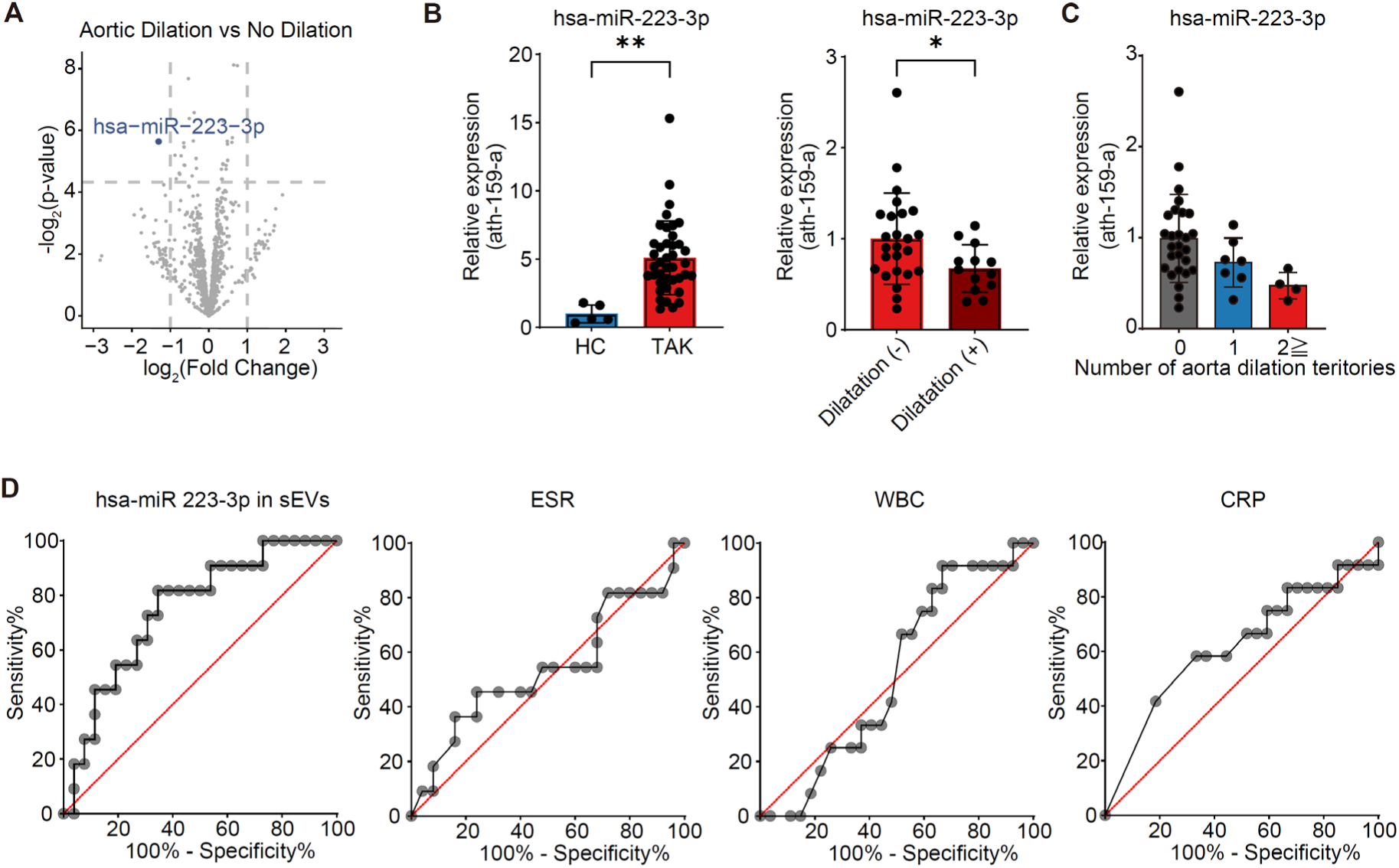
Identification of miR-223-3p as a potential biomarker for aortic dilation in patients with TAK (A) Volcano plot from microarray analysis comparing serum sEV-derived miRNA expression between patients with TAK having aortic dilation (n = 3) and those without aortic dilatation (n = 3). Significantly different expression was defined by a fold change > 2 and *p* < 0.05. (B) Quantitative PCR validation of miR-223-3p expression in serum sEVs. Expression levels were compared between HC and patients with TAK (left), and between patients with TAK, with and without aortic dilation (right). Expression values were normalized to the spike-in control ath-miR-159a (unpaired t-test). (C) miR-223-3p expression in patients with TAK stratified by the number of vascular dilation lesions (one-way ANOVA). (D) ROC curve analysis evaluating the predictive performance of serum sEV-derived miR-223-3p and conventional inflammatory markers. AUC, 95% confidence intervals calculated using Wilson score intervals, and p-values were determined for each marker. (B and C) Data are presented as mean ± SD. Abbreviations: ROC, receiver operating characteristic; AUC, area under the curve; WBC, white blood cell count; ESR, erythrocyte sedimentation rate; CRP, C-reactive protein; HC, healthy control; TAK, Takayasu arteritis; miRNA, microRNA; sEVs, small extracellular vesicles; PCR, polymerase chain reaction; ANOVA, analysis of variance.

To validate these results, we quantified serum sEV-derived miR-223-3p levels by qPCR. Compared to HC, patients with TAK generally exhibited increased miR-223-3p expression (Figure 4B, Left). However, within the TAK group, those with aortic dilation had significantly lower expression compared to those without dilation, confirming the microarray findings (Figure 4B, Right). Notably, miR-223-3p expression was not influenced by disease activity status or immunosuppressive therapy such as corticosteroids or disease-modifying antirheumatic drugs (Figure S4). These results suggest that the observed decrease in miR-223-3p expression is more closely related to aortic dilation and is unlikely to be solely explained by systemic inflammation or immunosuppressive therapy.

The 2022 ACR classification criteria for TAK consider the number of affected aortic and branch vessel territories as part of the disease definition^22^. Although not statistically significant, miR-223-3p expression in sEVs tended to decline as the number of aortic territories with dilation increased (Figure 4C), suggesting a potential association between miRNA levels and disease burden.

Finally, the diagnostic performance of miR-223-3p for identifying aortic dilation in patients with TAK was assessed by ROC curve analysis. Serum sEV-derived miR-223-3p archived an AUC of 0.7483 (95% confidence interval [CI]: 0.5822–0.9143; *p* = 0.0183), suggesting superior diagnostic potential compared to conventional markers including white blood cell (AUC = 0.5216; 95% CI: 0.3380–0.7053; *p* = 0.8313), ESR (AUC = 0.5418; 95% CI: 0.3198–0.7638; *p* = 0.6929), and CRP (AUC = 0.6188; 95% CI: 0.4150–0.8226; *p* = 0.2414) (Figure 4D).

Collectively, these findings suggest that decreased expression of sEV-derived miR-223-3p is closely associated with aortic dilation in patients with TAK and may serve as a promising non-invasive biomarker for vascular remodeling and disease progression.

### miR-223-3p Negatively Modulates NLRP3 Inflammasome Activation

To identify downstream targets of miR-223-3p encapsulated in circulating sEVs that differed according to the presence of aortic dilation in TAK, we employed TargetScan v8.0 to predict miRNA-target interactions^29^. Among the predicted targets, NLRP3 was identified as having a conserved binding site within its 3′ untranslated region (3′ UTR) across multiple mammalian species (Figure 5A).

**Figure 5.**
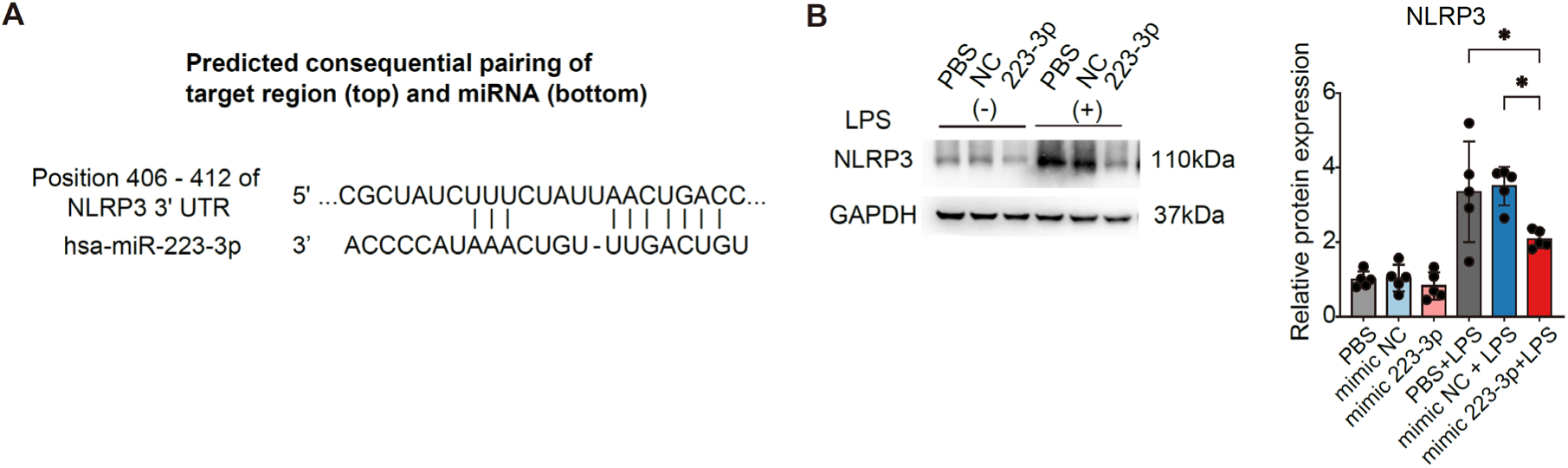
Suppressive role of miR-223-3p in NLRP3 activation (A) Predicted binding site of miR-223-3p in the 3′ UTR of NLRP3 mRNA, identified using TargetScan v8.0 and conserved across multiple mammalian species. (B) Western blot analysis of NLRP3 protein expression in THP-1-derived macrophages transfected with synthetic miR-223-3p mimics. Cells were differentiated with PMA and stimulated with LPS (One-way ANOVA). Data are presented as mean ± SD. **Abbreviations:** NLRP3, NOD-like receptor family pyrin domain containing 3; PMA, phorbol 12-myristate 13-acetate; LPS, lipopolysaccharide; UTR, untranslated region; SD, standard deviation

Given our previous finding that a missense mutation in max-like protein X (MLX), reported in patients with TAK, promotes NLRP3 inflammasome activation and contributes to disease pathogenesis^30^, we hypothesized that miR-223-3p may act as a negative regulator of NLRP3 expression at the post-transcriptional level. To test this hypothesis, THP-1 monocytes were transfected with synthetic miR-223-3p mimics or NC, differentiated into macrophages using PMA, and stimulated with LPS. Western blotting demonstrated that overexpression of miR-223-3p significantly suppressed NLRP3 protein expression, both under basal conditions and following LPS stimulation, compared to NC-transfected cells (Figure 5B). These findings indicate that miR-223-3p directly downregulates NLRP3, thereby attenuating inflammasome activation in macrophages.

### Identification of Potential Cellular Sources of Circulating sEVs Containing miR-223-3p in TAK

We investigated the potential cellular origins of miR-223-3p in circulating sEVs from patients

with TAK. miR-223-3p is abundant in sEVs released from activated platelets^31^ as well as in mononuclear cells and neutrophils^32^. Given that EVs retain surface markers and membrane proteins reflective of their cells of origin^33^, we characterized serum-derived EVs from patients with TAK by high-sensitivity flow cytometry. sEVs from both HC and patients with TAK exhibited a typical size distribution ranging from approximately 100 to 200 nm (Figure S5). Moreover, a substantial proportion of these vesicles expressed CD9, a well-established tetraspanin marker commonly used as a marker for sEVs (Figure 6A). We further evaluated the expression of representative surface markers corresponding to major blood cell types to further identify the cellular sources of circulating sEVs. Flow cytometric analysis revealed the presence of platelet-derived (CD41⁺), platelet- or megakaryocyte-derived (CD61⁺), and endothelial-derived (CD144⁺, CD31⁺) EVs (Figure 6B). In contrast, monocyte-derived (CD14⁺) and T cell–derived (CD3⁺) EVs were not detected in the serum of either patients with TAK or HC (Figure S5). Although the proportions of these marker-positive EVs did not differ significantly between patients with TAK and HC, the proportion and MFI of CD41⁺ EVs tended to be lower in patients with aortic dilatation compared to those without (Figures 6C, 6D).

**Figure 6.**
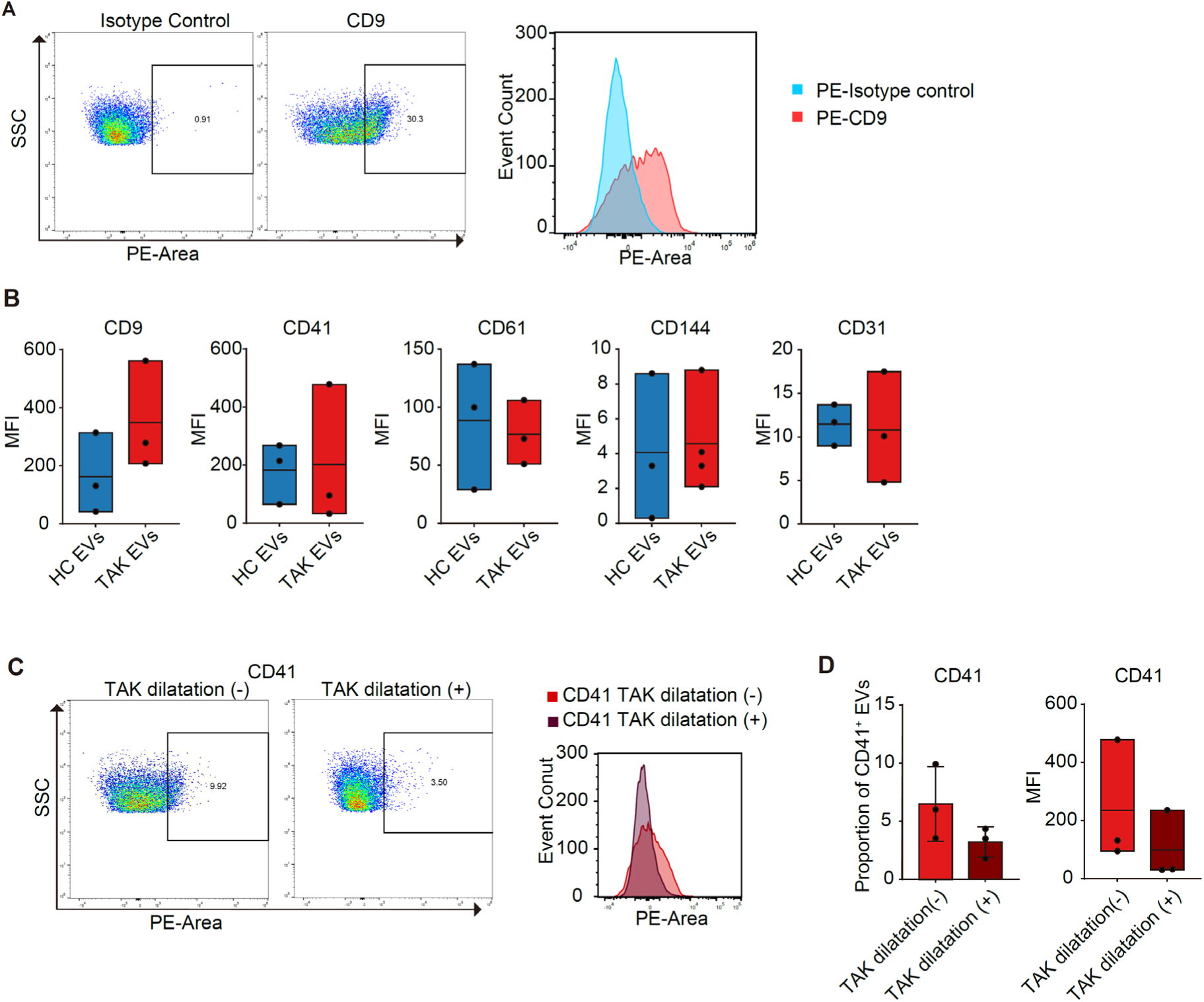
Identification of major cellular sources of circulating miR-223-3p–enriched sEVs in TAK (A) Flow cytometric characterization of serum-derived sEVs of isotype control (left) and CD9-stained EVs (middle). Overlaid histogram comparing CD9 and isotype control fluorescence (right). (B) Box plots comparing the MFI of EV surface markers (CD9, CD41, CD61, CD144, and CD31) between HC and TAK groups. (C) Representative flow cytometric dot plots of CD41⁺ EVs in patients with TAK, without (left) and with (middle) aortic dilation. The right panel shows the overlaid histogram of CD41 expression for both groups. (D) Quantitative comparison of CD41⁺ EVs between patients with TAK, with and without aortic dilation. The left graph displays the percentage of CD41-positive EVs; the right graph shows corresponding MFI values. (B and D) Box plots show means (lines), min to max (boxes). (D) Bar plots are presented as mean±SD. (B and D) (Mann–Whitney U test). **Abbreviations:** MFI, mean fluorescence intensity; SD, standard deviation; EV, extracellular vesicle; TAK, Takayasu arteritis

These results suggest that the reduction of circulating miR-223-3p observed in patients with TAK, with aortic dilatation, is largely attributable to a decreased number of platelet-derived EVs. Furthermore, disease-specific alterations in the selective packaging of miR-223-3p into EVs from different cell types may also contribute, highlighting an area for further investigation.

## Discussion

In this study, we investigated the role of sEVs in TAK pathogenesis and diagnosis. Our findings yielded three principal insights. First, circulating sEVs from patients with TAK exhibited altered miRNA profiles, with consistent downregulation of miR-223-3p in those with aortic dilation. Second, these sEVs promoted endothelial activation and monocyte adhesion in vitro, implicating their involvement in vascular inflammation. Third, target gene analysis revealed convergence on key immune regulatory pathways, particularly PTEN–PI3K–Akt signaling. Collectively, these findings highlight the potential of sEV-derived miRNAs as both biomarkers and modulators of vascular pathology in TAK.

Our in vitro experiments demonstrated that serum-derived sEVs from patients with TAK upregulated *ICAM-1* and *VCAM-1* expression in ECs and enhanced monocyte adhesion. In TAK, vascular inflammation is believed to originate in the vasa vasorum and the medio-adventitial junction, where mononuclear cell infiltration is frequently observed^13, 27^. Inflammation of large arteries typically involves recruitment of circulating monocytes and their differentiation into macrophages within the vessel wall^34–36^. Under normal conditions, large artery endothelium forms a tightly sealed barrier with minimal intercellular gaps, such that internalization by ECs represents the primary route through which circulating EVs deliver their cargo^37^. Collectively, our findings suggest that circulating sEVs may function as upstream mediators of endothelial activation, thereby linking systemic inflammation to local vascular pathology in TAK.

Bioinformatic analysis identified *PTEN* as a common predicted target of multiple miRNAs upregulated in TAK-EVs. Reduced *PTEN* expression in ECs has reportedly activated the PI3K-Akt signaling pathway and upregulated VCAM-1^38^ and ICAM-1^39^. Several of the upregulated miRNAs in our study—including miR-23b-3p, miR-26a-5p, and miR-92a-3p— have been experimentally validated to directly target and suppress *PTEN* expression^40–42^, and EV-delivered miRNAs reportedly modulate *PTEN* expression in recipient cells^43^. Enhanced PI3K-Akt pathway activation has also been described in the vascular endothelium of patients with TAK^44^. These observations raise the possibility that TAK-EV–derived miRNAs contribute to endothelial activation by modulating *PTEN* expression and PI3K-Akt signaling, although further experimental confirmation is required.

Among the differentially expressed miRNAs, miR-223-3p was uniquely downregulated in patients with TAK who had aortic dilation. Despite extensive investigation, no disease-specific biomarkers have been established for TAK, and clinical monitoring of disease activity still relies on non-specific inflammatory markers such as ESR and CRP^13, 22, 45^. Previously, we demonstrated that PTX3 can serve as a reliable disease activity marker independent of corticosteroid treatment, with elevated levels observed even in patients with relapsing TAK despite negative CRP^5^. Serum TNF-α and IL-6 levels correlate with disease activity^46^, while other candidates, including MMP3, MMP9, ICAM-1, and VCAM-1, have been suggested as potential markers^45^, although their clinical applicability remains uncertain. Our current findings suggest that EV-derived miR-223-3p may serve as a novel biomarker reflecting disease progression—specifically aortic dilation—independent of disease activity and immunosuppressive therapy. Given that routine clinical monitoring often requires repeated imaging with contrast-enhanced CT, PET-CT, or MRI, longitudinal assessment of exosomal miR-223-3p levels may offer a minimally invasive approach to complement imaging strategies. Prospective validation in larger cohorts will be essential to confirm its clinical utility.

Consistent with previous reports^47^, flow cytometric profiling of EV surface markers demonstrated that platelet-derived vesicles represent the predominant fraction of circulating sEVs in TAK. Although platelets are anucleate, they contain and process miRNAs^48^ and release EVs enriched in miRNAs, such as miR-223, upon activation^31^. Platelet miRNA cargo is dynamically regulated, as exemplified by miR-223-mediated autoregulation of P2RY12, which is impaired in type 2 diabetes due to defective Dicer processing^49^. In autoimmune and inflammatory disorders, such as rheumatoid arthritis, systemic lupus erythematosus, and vasculitis, platelet-derived microparticles have been reported to fluctuate with disease activity^50, 51^. In TAK, platelet indices have similarly been proposed as markers of disease activity^52, 53^, supporting a role for platelets in vascular pathology. Our findings suggest that platelet-derived EVs are a plausible source of circulating miR-223-3p in TAK, and that their abundance and miRNA cargo may be modulated by disease phase. Such temporal variability could influence EV miRNA signatures across different stages of TAK, emphasizing the need to account for disease phase when evaluating their potential utility as biomarkers.

Although this study provides important insights, it is subject to some limitations, and additional in vivo validation and analysis in larger patient cohorts are required to fully establish mechanistic links and biomarker utility.

In conclusion, our study identifies a distinct miRNA signature within circulating sEVs of patients with TAK and highlights miR-223-3p as a candidate biomarker of aortic dilation.

## Acknowledgments

We sincerely thank all the study participants for their involvement and also acknowledge the members of our laboratory for their valuable discussions and support pertaining to this work.

## Supplemental Material

Table S1 Figure S1-S5

Major Resources Table

## Non-standard Abbreviations and Acronyms

TAK: Takayasu arteritis
HC: Healthy controls
EV: Extracellular vesicle
sEV: Small extracellular vesicle
TAK-EV: sEV derived from patients with TAK
HC-EV: sEV derived from healthy controls
miRNA: MicroRNA
SEC: Size-exclusion chromatography
ECs: endothelial cells
ICAM-1: Intercellular adhesion molecule-1
VCAM-1: Vascular cell adhesion molecule-1
PTEN: phosphatase and tensin homolog NLRP3 NLR family pyrin domain containing 3

## Novelty and Significance

### What Is Known?

- Takayasu arteritis (TAK) is a large-vessel vasculitis in which chronic vascular inflammation can lead to stenosis or aortic dilatation, however, circulating biomarkers for vascular remodeling in TAK are limited.
- Extracellular vesicles (EVs) carry molecular cargo reflective of their cellular origin and activation state, and EV-derived microRNAs (miRNAs) have emerged as potential biomarkers in other cardiovascular diseases.

What New Information Does This Article Contribute?

- Provides the first integrated characterization of circulating small EVs in TAK, combining high-sensitivity flow cytometry with miRNA profiling and pathway analysis.
- Patients with TAK exhibit distinct circulating sEV miRNA signatures linked to PI3K– Akt signaling, and platelet-derived EV–associated miR-223-3p is selectively downregulated in those with aortic dilation.
- miR-223-3p outperforms CRP and ESR in discriminating aortic dilatation, highlighting its potential as a biomarker of vascular remodeling.

## Paragraph Summary

Takayasu arteritis (TAK) is an inflammatory arteriopathy in which progressive vascular remodeling can result in aortic dilatation, yet reliable circulating biomarkers for this phenotype are lacking. This study provides the first comprehensive characterization of circulating small extracellular vesicles (sEVs) in TAK, integrating high-sensitivity flow cytometric phenotyping with miRNA profiling and pathway enrichment analysis.

Functionally, TAK-EVs promoted endothelial activation and monocyte adhesion in vitro, and their miRNA cargo was enriched for PI3K–Akt pathway regulators, with *PTEN* identified as a common predicted target. Comparative analysis revealed that platelet-derived EV– associated miR-223-3p was uniquely downregulated in patients with aortic dilatation, independent of disease activity or immunosuppressive therapy. AUC demonstrated that miR-223-3p discriminates aortic dilation more effectively than conventional inflammatory markers such as CRP and ESR. These findings suggest that EV-derived miR-223-3p may serve as a mechanistically relevant and clinically useful biomarker of vascular remodeling in TAK, linking platelet biology to disease progression.

